# LC-SRM combined with machine learning enables fast identification and quantification of bacterial pathogens in urinary tract infections

**DOI:** 10.1101/2024.05.31.596829

**Authors:** Clarisse Gotti, Florence Roux-Dalvai, Ève Bérubé, Antoine Lacombe-Rastoll, Mickaël Leclercq, Cristina C. Jacob, Maurice Boissinot, Claudia Martins, Neloni R. Wijeratne, Michel G. Bergeron, Arnaud Droit

## Abstract

Urinary tract infections (UTIs) are a worldwide health problem. Fast and accurate detection of bacterial infection is essential to provide appropriate antibiotherapy to patients and to avoid the emergence of drug-resistant pathogens. While the gold standard requires 24h to 48h of bacteria culture prior MALDI-TOF species identification, we propose a culture-free workflow, enabling a bacterial identification and quantification in less than 4 hours using 1mL of urine. After a rapid and automatable sample preparation, a signature of 82 bacterial peptides, defined by machine learning, was monitored in LC-MS, to distinguish the 15 species causing 84% of the UTIs. The combination of the sensitivity of the SRM mode on a triple quadrupole TSQ Altis instrument and the robustness of capillary flow enabled us to analyze up to 75 samples per day, with 99.2% accuracy on bacterial inoculations of healthy urines. We have also shown our method can be used to quantify the spread of the infection, from 8×10^4^ to 3×10^7^ CFU/mL. Finally, the workflow was validated on 45 inoculated urines and on 84 UTI-positive urine from patients, with respectively 93.3% and 87.1% of agreement with the culture-MALDI procedure at a level above 1×10^5^ CFU/mL corresponding to an infection requiring antibiotherapy.

**HIGHLIGHTS:**

– LC-MS-SRM and machine learning to identify and quantify bacterial species of UTI
– Fast sample preparation without bacterial culture and high-throughput MS analysis
– Accurate quantification through calibration curves for 15 species of UTIs
– Validation on inoculations (93% accuracy) and on patients specimens (87% accuracy)

## INTRODUCTION

Urinary tract infections (UTIs) are one of the most common bacterial infections. 400 million people in the world contracted a UTI in 2019 and more than 230,000 of them died from it (1). UTIs mainly affect women (50-60%) and it has been shown that half of them will have at least one in their life, reducing their quality of life (2). UTIs are therefore a global health problem, with an estimated economic impact of $3.5 billion per year in the United States alone (3).

To date, the gold standard for detection and identification of bacterial cells in urine consists in a microbiological culture, followed by bacterial species identification using MALDI-TOF (matrix-assisted laser desorption ionization–time of flight) mass spectrometry analysis (4–6). Due to the length of this culture, results can only be obtained in 24 to 48h (1, 7), delaying appropriate antimicrobial treatment. Moreover, during this time, broad-spectrum antibiotics are often administered to the patient empirically, although this practice has been shown to be responsible for the occurrence of antimicrobial-resistant pathogens in the population (8–10). To preserve our therapeutic arsenal and prevent the emergence of multidrug resistant pathogens, considerable efforts must be made to shorten the analysis time required to identify the pathogens responsible for infections, particularly in the case of UTIs. Rapid detection of as low as 1×10^5^ colony forming units per mL of urine (CFU/mL), the clinical threshold at which the urinary tract is considered infected, remains the main analytical challenge (11).

In the past few years, several studies have investigated culture-free methods to reduce this time to less than 4 hours (12–15). For example, Veron L. *et al.* (14) proposed three alternative methods of urine preparation (differential centrifugation, filtration or short bacterial culture) prior to MALDI-TOF analysis. They showed that better results were obtained with a 5h culture step. Similar results have been obtained by Lei D. *et al.* (15) and also propose a short term culture to enrich bacterial colonies before the MALDI-TOF analysis. Although these studies show an improvement of turnaround time to obtain an identification, the lack of sensitivity of the MALDI-TOF analysis still requires an enrichment of bacterial cells.

Another study (16) has proposed a new method called FLAT (Fast Lipid Analysis Technique) in which urine or bacterial pellets are directly spotted on the MALDI plate prior to lipids extraction and MALDI-TOF analysis. This very fast (< 1 hour) and simplified (minimal hand-on time) assay allows to reach 94% of concordance with the gold-standard culture procedure but remains limited to the identification of Gram-negative species, which are predominantly responsible for UTIs (3, 17). Moreover, no quantification of bacteriuria can be obtained with these strategies, while it is essential to distinguish urine contamination (1×10^3^ to 1×10^4^ CFU/mL) from infection requiring antibiotherapy (> 1×10^5^ CFU/mL), and thus avoid unnecessary treatment (18). As for the gold standard method combining bacterial culture and MALDI-TOF, an approximate range of bacteriuria is given through a visual estimate of the number of colonies on the culture plate based on the technician’s expertise.

Other methods, not based on MALDI-TOF analysis, have been proposed. Polymerase Chain Reaction (PCR) for instance amplifies specific regions of the genome such as highly conserved 16S ribosomal subunit RNA gene, often limiting the identification of the bacteria at the genus level (19). Although efforts have been made in 2019 by Johnson *et al.* (20) to give resolution at the species and even at the strain level by considering full-length 16S intragenomic copy variants, this technique requires the use of primers and thus can only detect a limited number of predefined species (21). These limitations could be addressed with the whole genome sequencing (WGS) method, which enables high-throughput and untargeted genome sequencing. However, its high cost, the computational infrastructure and the time required to obtain a result limit its deployment in clinical laboratories (22). In contrast, Liquid Chromatography - tandem Mass Spectrometry (LC-MS/MS) appears to be a promising method for diagnosing UTIs. Thanks to its sensitivity and specificity, this method has replaced MALDI-TOF mass spectrometry in most research applications. Moreover, its low cost and automation capability make it suitable for routine use.

In bottom-up proteomics by LC-MS/MS (23, 24), proteins are extracted from various biological matrices and digested with a trypsin enzyme. The resulting peptides are then separated based on their hydrophobicity on a reverse phase chromatography column allowing the reduction of the sample complexity to be analyzed by the mass spectrometer (25). In contrast to MALDI, for which a limited number of mass spectra can be obtained, this strategy enables the acquisition of thousands of them, therefore increasing the specificity. Moreover, the signal intensity is correlated to the amount of peptide present in the sample allowing quantification values to be obtained (26, 27). Nowadays, very sensitive and high-throughput LC-MS/MS systems enable the detection and the quantification of low abundance molecules among complex matrices such as urine (28). In 2019, our team used LC-MS/MS to develop a new method to detect bacteria directly from urine samples without the need for bacterial culture (29). To do so, we trained machine learning algorithms with bacterial peptide identifications obtained by LC-MS/MS. Then, we defined a list of 82 peptides (UTI signature) able to distinguish 15 bacterial species causing 84% of the UTIs. By monitoring the UTI signature in targeted analyses, we demonstrated the method can correctly predict 97% of the bacterial species inoculated in healthy urine in a range from 2.5×10^4^ to 8.8×10^6^ CFU/mL. The prediction accuracy reached 100% for inoculation > 1×10^5^ CFU/mL. All the samples were injected into a nanoflow liquid chromatography with a 90 minutes gradient, interfaced to a high resolution Orbitrap Fusion mass spectrometer, widely used in the proteomic research field. While very sensitive analysis could be achieved with nano LC-MS/MS, its lack of robustness prevents its use in routine analysis. Furthermore, the associated costs for the acquisition and maintenance of high resolution instruments are not suited to clinical laboratories. In addition, with 90 min gradients, less than 12 samples could be analyzed every day, which is not in line with clinical throughput. Over the past decade, LC-MS/MS in selected / multiple reaction monitoring (SRM / MRM) has emerged as a powerful approach in clinical proteomics (30–32). Typically performed on affordable triple quadrupole instruments, SRM is characterized by its high selectivity, sensitivity and reproducibility (33). This acquisition mode consists in monitoring the precursor / fragment transitions from a peptide of interest, avoiding signal interference from small molecules or other proteins present in the sample matrix. Moreover, as only a limited number of transitions (hundreds) is usually monitored in a single run, capillary or micro-flow LC can be used, increasing the speed and robustness of the analysis (34, 35).

In this work, we aimed to improve our previously published strategy for the identification of bacterial species in urine by transferring the monitoring of UTI signature from high resolution to triple quadrupole instruments. This new workflow, called LC-SRM-ML, combines the strengths of capillary flow LC and SRM acquisition to make it suitable for routine analysis in clinical laboratories. Furthermore, we defined calibration curves of the MS signal to enable not only the identification but also the quantification of the 15 bacterial species most frequently found in UTIs, in less than 4 hours without bacterial culture.

## EXPERIMENTAL PROCEDURES

### Experimental design and statistical rationale

For the first part of this study, which consisted in transferring the method from an Orbitrap to a triple quadrupole instrument, 15 bacterial species were spiked in 4 different urines from healthy volunteers at 6 concentrations. This dataset, called *Transfer - Inoculations*, was used to train a new machine learning model and to generate calibration curves.

The reproducibility (*Transfer - Reproducibility*) of the workflow was evaluated by inoculating 4 bacterial species (Eco, Efa, Kpn and Sag) in 50 mM Tris or urine backgrounds, in 6 replicates. The technical reproducibility from sample preparation to the LC-SRM analysis was evaluated with the Tris background. The analytical (injection) reproducibility was evaluated by injecting a pool of inoculations prepared in the 50 mM Tris background 6 times. Finally, to estimate the biological reproducibility (inter-individual variation), uropathogens were spiked into urines collected from healthy volunteers or from UTI-negative patients, with an optional rapid centrifugation (1 min at 100x*g*) at the beginning of the sample preparation, to remove large cell debris.

The LC-SRM-ML workflow was validated on 45 inoculated samples (15 bacterial species inoculated at 3 concentrations in urines from healthy volunteers) (*Validation - Inoculations*) and with 84 UTI-positive urines from patients (*Validation - Patients*) collected at the CHU de Québec. All the details of the 4 datasets could be found in the Supplemental Table S1.

Samples from the *Transfer - Inoculations* dataset were injected from low to high concentrations, whereas validation samples (inoculations and patients) were injected in a randomized order. Intensity tables from Skyline software exports were used to discretize the data used as input for ML. They were also used to generate linear regressions and to calculate Pearson correlation and relative standard deviation (RSD) using the R software (36). The statistics are described in the relevant following sections.

### Bacterial culture

Bacterial strains were obtained from the Culture Collection of Centre de Recherche en Infectiologie of Université Laval (CCRI, Québec, Canada) registered as WDCM861 at the World Data Centre for Microorganisms. The bacterial strains used and their corresponding culture conditions are listed in Supplemental Table S2. Each bacterial strain was plated on blood agar (TSA II 5% Sheep blood; Becton Dickinson) and incubated at 35°C overnight under aerobic conditions (or in air + 5% CO_2_ for *S. agalactiae* and *S. mitis*). Bacterial colonies were then collected and resuspended in phosphate-buffered saline (PBS, 137 mM NaCl / 6.4 mM Na_2_HPO_4_ / 2.7 mM KCl / 0.88 mM KH_2_PO_4_ at pH 7.4) at an optical density of 0.5 MacFarland. 30 µL of this suspension were used to inoculate 3 mL of BHI medium (Brain Heart Infusion medium broth; Becton Dickinson) which was incubated overnight at 37°C under aerobic conditions with agitation (or 35°C in air + 5% CO_2_ without agitation for *S. agalactiae* and *S. mitis*). Finally, broth cultures were done by inoculation of 30 µL of the previous culture in 3 mL BHI medium and incubated in the same conditions. The cultures were stopped in an exponential phase according to the previously known semi-log growth curve of each strain (Supplemental Table S2). Bacterial cells in these cultures were counted in triplicates by incubation of 50 µL of serial dilutions on Blood agar plates, at 35°C overnight under aerobic conditions (or in air + 5% CO_2_ for *S. agalactiae* and *S. mitis*).

### Urine collection and bacteria inoculations

This study is classified as a Tier 3 level. Urine from healthy volunteers or from patients (UTI-negative or positive) were collected at the CHU de Québec - Université Laval hospital, according to an ethical protocol approved by the Comité d’Éthique de la Recherche du CHU de Québec - Université Laval (recording number 2106-2656). Only midstream urines, from adults with a minimum volume of 10 mL were included. Collected urines were kept at 4°C during a maximum of 2 days before sample preparation with the LC-SRM-ML workflow. The urine specimens were anonymized and only the MALDI-TOF analysis reports (identification of the species and the quantification) was transmitted to the research group.

The information of uropathogens inoculations (concentrations, replicates) can be found in the Supplemental Table S1.

### Standard workflow (culture and MALDI-TOF)

For the culture-MALDI workflow used in *Validation - Inoculations* and *Patients* datasets, the standard procedure of the CHU de Québec - Université Laval hospital microbiology laboratory was used. Briefly, 1 µL of urine was plated on blood agar and incubated at 35°C for 18h to 24h, under aerobic conditions. The number of colonies was estimated (0 colonie: < 1×10^3^ CFU/mL; 1-9 colonies: 1×10^3^ - 1×10^4^ CFU/mL; 10-99 colonies: 1×10^4^ - 1×10^5^ CFU/mL and > 100 colonies: > 1×10^5^ CFU/mL) to evaluate the level of the infection. Then, a colony is isolated and spotted on a MALDI plate with 1 µL of α-Cyano-4-Hydroxycinnamic Acid matrix. MALDI-TOF/MS analysis was performed on a Vitek® MS instrument (bioMerieux).

### LC-SRM-ML workflow

#### Sample preparation

Samples from inoculations in healthy urines or from patient urine specimens were processed and analyzed in the same way, with the exception of a rapid centrifugation (1 min at 100x*g*) that was added at the beginning of the process for patient specimens, to remove large human cell debris. Bacterial cells were then isolated by centrifugation at 10,000x*g* during 10 min. Pellet was washed with 1 mL of 50 mM Tris and centrifuged again for 10 min at 10,000x*g*. The supernatant was discarded and the pellet was frozen on dry ice for 5 min. Cell lysis was performed by resuspension of the pellet in 50 mM ammonium bicarbonate containing 20 units of mutanolysin enzyme (Mutanolysin from *Streptomyces globisporus* ATCC 21553, Sigma-Aldrich) and incubation at 37°C for 1h. Bacterial cells inactivation and protein denaturation was then achieved by addition of 1% sodium deoxycholate (SDC) and 20 mM DTT (dithiothreitol) for 10 min and heating at 95°C. Proteins were then digested using 250 ng of trypsin enzyme (Sequencing Grade Modified Trypsin, Promega) and incubation for 1h at 37°C. Samples were acidified with 5 µL of 50% formic acid (FA) to stop the digestion and precipitate the SDC. After 10 min of centrifugation at 10,000x*g*, peptides contained in the supernatant were purified on C_18_ StageTips (37), vacuum-dried and stored at −30°C until mass spectrometry analysis.

#### Mass spectrometry analysis

For each sample, peptides were resuspended in 10.5 µL of 0.1% FA containing 40 fmol of Dionex^TM^ Cytochrome C digest (Life Technologies) (CytoC) prior to LC-SRM injection. 5 µL of each sample was injected on a Vanquish Neo (Thermo Fisher Scientific) liquid chromatography system, in trap-and-elute mode. Peptides were trapped on a PepMap™ Neo Trap column (Thermo Fisher Scientific, 300 μm x 5 mm, 5 μm particles, 100 Å pore) at the maximum of 60 μL/min or 500 bar. Then, the pre-column was switched on-line with a 50 cm nanoμPAC column (Thermo Fisher Scientific, heated at 60°C) and the peptides were eluted from 5% to 50% of solvent B (A: 0.1% FA; B: 80% ACN / 0.1% FA) in 10 min, at a 2 µL/min flow-rate. Column ending was connected to an EASY-spray source with a 20 µm silica emitter (Pepsep) and interfaced with a TSQ Altis mass spectrometer (Thermo Fisher Scientific). Spray voltage was set to 1900V and the ion transfer tube was heated at 275°C.

#### Development and Analytical Validation Targeted MS Assays/Measurements

The SRM assay was developed on a TSQ Altis instrument operating with the Tune version 3.4.3279 software. In brief, the 6 most intense fragments of the DDA spectral library from our previous study were used as initial transitions (29). Collision energy (CE) was optimized on each of the precursor/fragment transitions using Skyline, with the TSQ Quantiva CE formula (calculated as CE = 0.0339 m/z + 2.3597 or CE = 0.0295 m/z + 1.5123, respectively for doubly or triply charged ions). A ramping of 11 CE values (+/- 5 around the initial CE) of 2 voltages was performed. In the optimized method, the 4 most intense precursor/fragment transitions along with their best CE were retained for each peptide for a total of 352 transitions (82 peptides from UTI signature + 6 peptides from CytoC standard). Transition priorisation function of the Tune software was used to increase the dwell time for the transitions used for peptide quantification (Supplemental Table S3).

#### Data processing

Peptide peak detection was performed using Skyline software (version 23.1.0.268). To determine the retention times of all the UTI signature peptides and distinguish them from background noise, we used the fact that bacterial peptide intensity should increase with the 6 inoculated concentrations in the *Transfer - Inoculation* dataset. An iterative process was finally performed by manually curating the peptide peaks over the 24 samples of each of the 15 bacterial species to confirm the retention times (Supplemental Table S4). A peptide was considered as detectable if at least 3 of the 4 transitions overlapped and if the S/N ratio was > 3.

In the 3 other datasets (*Transfer DIA to SRM - Reproducibility, Validation - Inoculations, Validation - Patients*), peptide peaks were unselected in Skyline if not present at their expected retention time or reintegrated if necessary. For some urines, a retention time shift could be observed for all the peptides of the signature and those of the CytoC standards, probably due to an effect of the biological matrix on the column. In that case and if there is no ambiguity on the detected peptide, peaks were considered anyway. Areas under the chromatographic peak from each transition were then exported from Skyline in a .csv format to be processed in R software.

##### Quality control and normalization of analyzed samples

The signals of the 6 CytoC standards peptides spiked in each sample were used to check the quality of the injections. The log_2_ of the median signal of the 6 best CytoC transitions was calculated for each sample and used to calculate the standard deviation across the 360 samples of the *Transfer - Inoculations* dataset. Then, a sample was considered as compliant if its log_2_ CytoC median signal fell in the range of +/- 3 standard deviation calculated previously (Supplemental Figure S1). Moreover, to calculate the bacterial concentration in a sample based on the SRM signal, the best transitions for each peptide were kept and were normalized by the previously calculated median.

##### Machine learning training

The list of detected peptides in each sample was first discretized as TRUE when the peptide was present, and FALSE if not. The 360 inoculations (15 bacteria x 6 concentrations x 4 replicates) from the *Transfer - Inoculations* dataset were used to perform a machine learning training. An additional 15 samples were artificially annotated as FALSE for all the 82 peptides and added to this dataset, for a total of 375 samples, in order to predict a class called “*Unknown*” when not enough peptides are present to correctly predict a bacteria from the list of the 15. The machine learning training was handled with BioDiscML (version 1.8.14) (38), a sequential minimal optimization algorithm for machine learning based on the Weka Java library (39). This tool is able to train thousands of models employing various classifiers with many hyperparameters combinations. Thus, a variety of machine learning models have been used for classification of bacteria, including rules-based methods (*i.e.*, OneR, PART, Prism, Ridor, and RoughSet), decision tree models (*i.e.*, BFTree, J48, RandomForest, and DecisionStump), Bayesian models (*i.e.*, NaiveBayes and BayesNet), lazy classifiers (*i.e.*, KStar and LocalKnn), and function-based models (*i.e.*, Logistic, SimpleLogistic), Multilayer Perceptron and support vector machine, among others. To avoid overfitting, an automated stratified sampling is performed to create a validation set and all models undergo evaluation through cross-validation methods like k-fold, Bootstrapping, and repeated holdout, where standard deviation is also determined. In our case, the random sampling was split in 2/3 (248 samples) and 1/3 (127 samples) as train and test sets. Then, from the many models generated, we selected the best model based on the global predictive performance revealed by average Matthews Correlation Coefficient (MCC) based on several evaluations (10-fold cross-validation, leave-one-out-cross-validation, holdout, repeated holdout, and bootstrapping) and small number of interactions.

##### Machine learning prediction

To predict the bacterial species in new samples, a list of discretized peptides (TRUE/FALSE) were provided either to the first generation model based on DIA data or to the best model identified with SRM data. Only samples having at least 2 bacterial peptides are used for prediction, else they were labeled as “*Non-infected*”.

##### Quantification - Generation of calibration curves

The data from *Transfer - Inoculations* dataset were used to obtain calibration curves for each bacterial species: for each peptide detected, the best transition (priority 1 in Supplemental Table S3) was kept and was normalized by the median of peak area of the 6 best transitions of the CytoC standard. Then, the normalized area from each transition was averaged across the 4 biological replicates (Supplemental Table S5). Linear regressions were calculated between the log_2_(Area) and the log_10_ of inoculated bacterial concentration (CFU/mL of urine). To quantify a bacterial species, only 3 peptides, called “*quantifier peptide*”, were selected on the following criteria: detectable in at least 85% of the samples (21 out of the 24 inoculations), with a coefficient of determination *r^2^*> 0.970 and with the highest intensities (Supplemental Table S6). Visual inspection of the peptides was also carried out in Skyline to avoid picking peptides affected by background interference signals.

Pearson correlation coefficients were calculated for the 3 quantifier peptides for each species, using between the log_2_(Area) of each replicate independently and the log_10_ of bacterial concentration inoculated (CFU/mL of urine) (Supplemental Table S7).

##### Quantification - Calculation of the bacterial concentration in urine of unknown samples

Only the 3 quantifier peptides for the detected bacteria were retained and the concentration was calculated as concentration (CFU/mL) = (Area - intercept) / slope. Finally, the average of the concentrations calculated for the 3 transitions was performed in order to estimate the bacterial quantification of the sample.

##### Quantification - Evaluation of the reproducibility

The normalized signal of the 3 quantifier peptides of each species was used to calculate relative standard deviation (RSD) from the mean, over the 6 replicates of each type of reproducibility to be evaluated (analytical: inoculations prepared in 50 mM Tris, pooled just before injection and injected 6 times; technical: 6 independently prepared inoculations in 50 mM Tris; biological: inoculations prepared in urines from healthy donors, from UTI-negative patients or from centrifuged (1 min at 100x*g*) UTI-negative patients). The RSD (%) of each quantifier peptide was calculated as RSD (%) = standard deviation across the 6 replicates / average of the normalized area of the 6 replicates x 100 (Supplemental Table S8). In the case where a peptide is not detected in all 6 replicates of a condition, the RSD is calculated on the others, with at least a minimum of 4 replicates.

##### Validation on inoculated urines - Pearson correlation

Pearson correlation coefficient was calculated between the bacterial concentration inoculated in healthy urine (based on broth culture plate counting) and the concentration calculated with the LC-SRM-ML method (based on linear regressions).

## RESULTS

### Analysis workflow

In this work, we propose an analytical workflow to reduce the turnaround time to obtain the identification of the bacterial species responsible for a UTI. Figure 1 shows the entire workflow comprising bacterial isolation by centrifugation, protein extraction and trypsin digestion, monitoring of machine learning-defined UTI peptide signature by LC-MS in SRM mode and bioinformatics processing. The whole procedure takes less than 4 hours.

**Figure 1:**
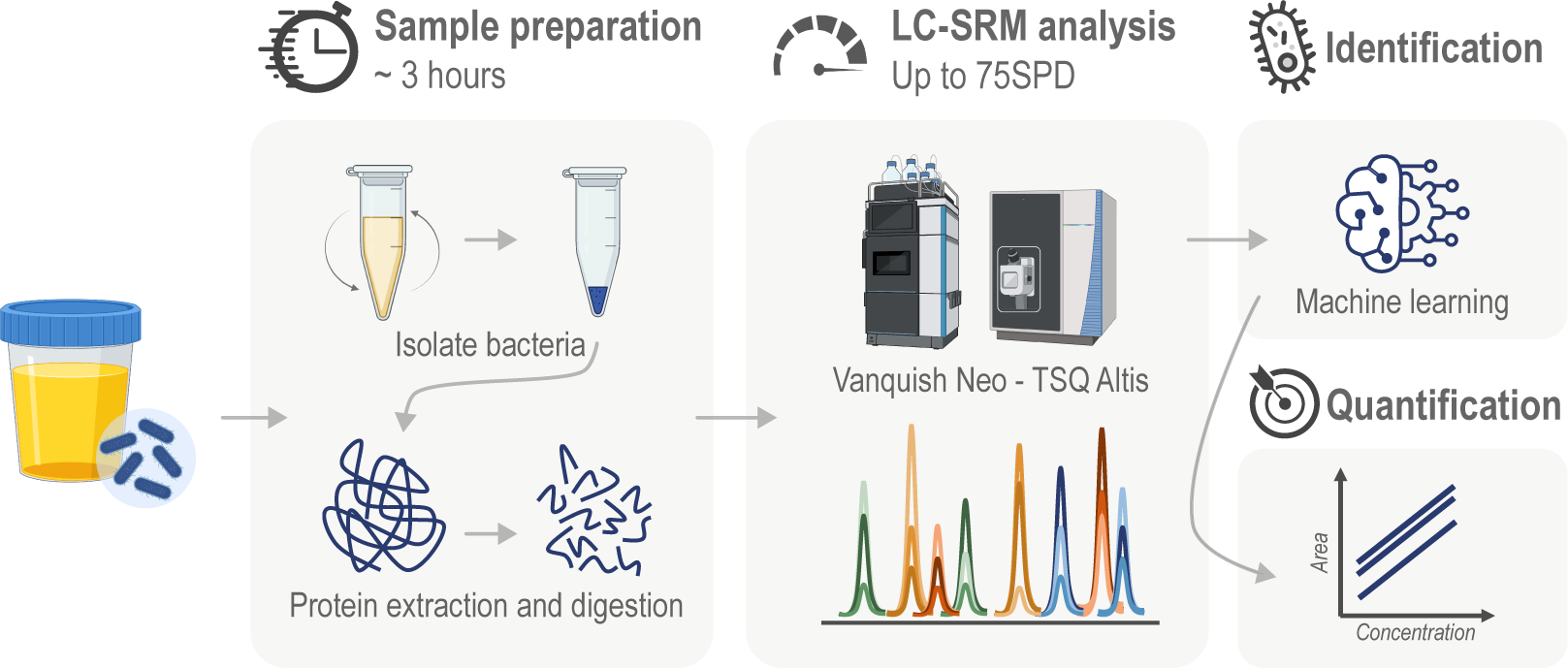
LC-SRM-ML analysis workflow. The workflow is composed of 4 steps: the “sample preparation” to isolate bacterial cells from urine, extract and digest their proteins into peptides; the “LC-SRM analysis” to monitor peptides of UTI signature; the “identification” to predict bacterial species using a machine learning model and the “quantification” to estimate bacterial amount in urine using SRM peptide signal with standard curves.

Compared to our previously published work (29), we improved the method by reducing the initial volume of urine required from 10 mL to just 1 mL, enabling higher throughput during sample preparation as bacterial isolation can be performed with benchtop centrifuges. All the following steps of bacterial lysis and protein digestion can be performed in less than 3 hours and is entirely automatable. Peptides are then analyzed with a robust LC-SRM method. The gradient length was reduced to 10 min with a microflow rate (2 µL/min) allowing analysis of up to 75 samples per day. 352 transitions (from the 82 UTI peptide signature and 6 CytoC standard peptides) are monitored throughout the LC gradient, thus preventing any loss of signal due to a possible shift in retention time that could be caused by a biological matrix effect on the column. The list of detected peptides is then sent to a prediction model derived from machine learning to predict the bacterial species causing the infection. Once the identification has been made, quantification can be obtained by measuring the area under the chromatographic peaks of 3 different peptides.

### Monitoring of the UTI signature with triple quadrupole instruments

#### Identification by LC-SRM-ML

As the UTI signature has been developed on high resolution Orbitrap instruments (29), we first aimed to show our LC-MS-ML workflow could be transferable to a robust triple quadrupole mass spectrometer. To do so, we used inoculations of the 15 species of interest in urines of 4 healthy volunteers (biological replicates) at 6 concentrations, in order to mimic the infection found in UTIs (Supplemental Table S1).

Over the 360 inoculated samples, 90.3% were accurately predicted with the first generation model based on DIA data (Supplemental Figure S2A), corresponding to an AUC = 0.996. 6.4% were misclassified and 3.3% were considered as non-infected (“*Blank*“). For the samples predicted as non-infected, multiple bacterial peptides were detected but not enough for the model to correctly classify them. Most of the error of the ML model comes from *K. aerogenes* which is predicted as *K.pneumoniae* in 85% of the cases. These two species are closely related (same genus) and share 21 peptides of the UTI signature. When we compare peptides of the UTI signature detected with the Orbitrap Fusion in DIA mode and with the TSQ Altis in SRM mode for these two bacteria (Supplemental Figure S2B), we can notice that 5 peptides were totally discriminant in DIA (respectively 3 and 2 peptides specific for *K. pneumoniae* and *K. aerogenes*) which was not the case with the same samples analyzed in SRM. No specific peptide was detected in SRM for *K. aerogenes*, leading to misclassification of this bacteria with *K. pneumoniae*, probably due to a different ionization of these peptides on the two instruments.

To improve the quality of prediction, two modifications were added to the initial method. First, to avoid the prediction of urine specimens as “Blank” (*i.e* non-infected) when the bacterial signal is present but too low to obtain an identification, we considered the urine samples as infected if at least 2 bacterial peptides of the UTI signature were detectable. Then, the same SRM data from 360 inoculations as well as 15 artificial data called “*Unknown samples*”, added to mimic non-infected urines, were used to train a new model to prevent confusion with *K. aerogenes* and *K. pneumoniae*. The best identified model was a Random Forest model with 10 trees. Two third of the dataset (248 samples) were used as a training set and reported 98.4% of correct classifications using a 10 fold cross validation, the remaining third (127 inoculated samples) were used as a test set for which only 1 inoculation was incorrectly predicted (*S. aureus* predicted as *S. haemolyticus*), corresponding to a 99.2% accuracy (Figure 2).

**Figure 2:**
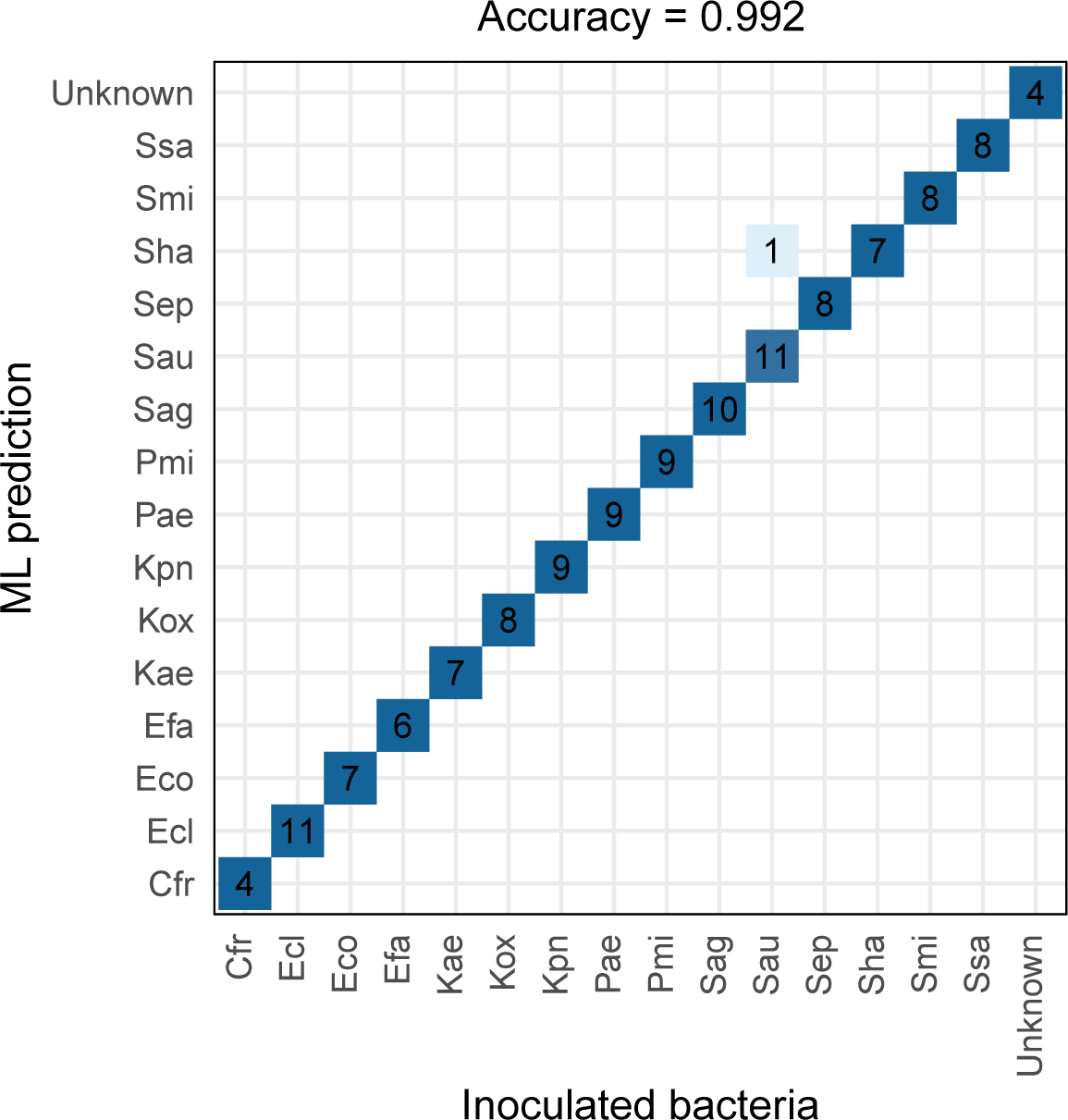
Bacterial identification by machine learning on inoculated urines. The 82 peptides of the UTI signature were monitored in LC-SRM in 360 samples (24 inoculations for each of the 15 bacterial species) and 2/3 were used to train a new random forest model. The confusion matrix reports the number of samples predicted by ML over the 127 samples of the test set.

#### Quantification using linear regressions

After the bacterial species had been identified by the UTI ML model, areas under the curve of the peptide chromatographic peaks were extracted from Skyline software and used to generate quantitative data for the samples considered as infected. The log_2_ of area and the log_10_ of the bacterial concentration in urine (CFU/mL) were used to calculate linear regressions for each peptide of each species (Supplemental Figure S3 and Supplemental Table S5). Finally, 3 peptides (further named “quantifier peptides”) were manually selected for each species to quantify them, based on the following criteria: detectable in at least 85% of the samples (21 inoculations over the 24), having a coefficient of determination *r^2^* > 0.97 and being among the highest intensities (Supplemental Table S6). A visual inspection of the peptides was also performed in Skyline software to avoid taking peptides affected by background signal interferences. Figure 3A shows linear regressions of the 3 peptides selected for the quantification of each bacteria. The average of the coefficient of determination (*r^2^*) values across the 45 peptides was 0.994 (Figure 3B; Supplemental Table S7). The reproducibility of the quantification for each peptide across the 4 biological replicates was evaluated with the Pearson correlation factors, which were 0.948 in average. Moreover, the signal to noise ratio (S/N) calculated at the lowest concentration was ranging from 4 to 188, with an average of 41. Thus, we were considering that concentration as our lower limit of quantification (LLOQ) (40–42). All these metrics suggest our method could be used for quantification, in a wide range of concentrations, representing 3 orders of magnitude.

**Figure 3:**
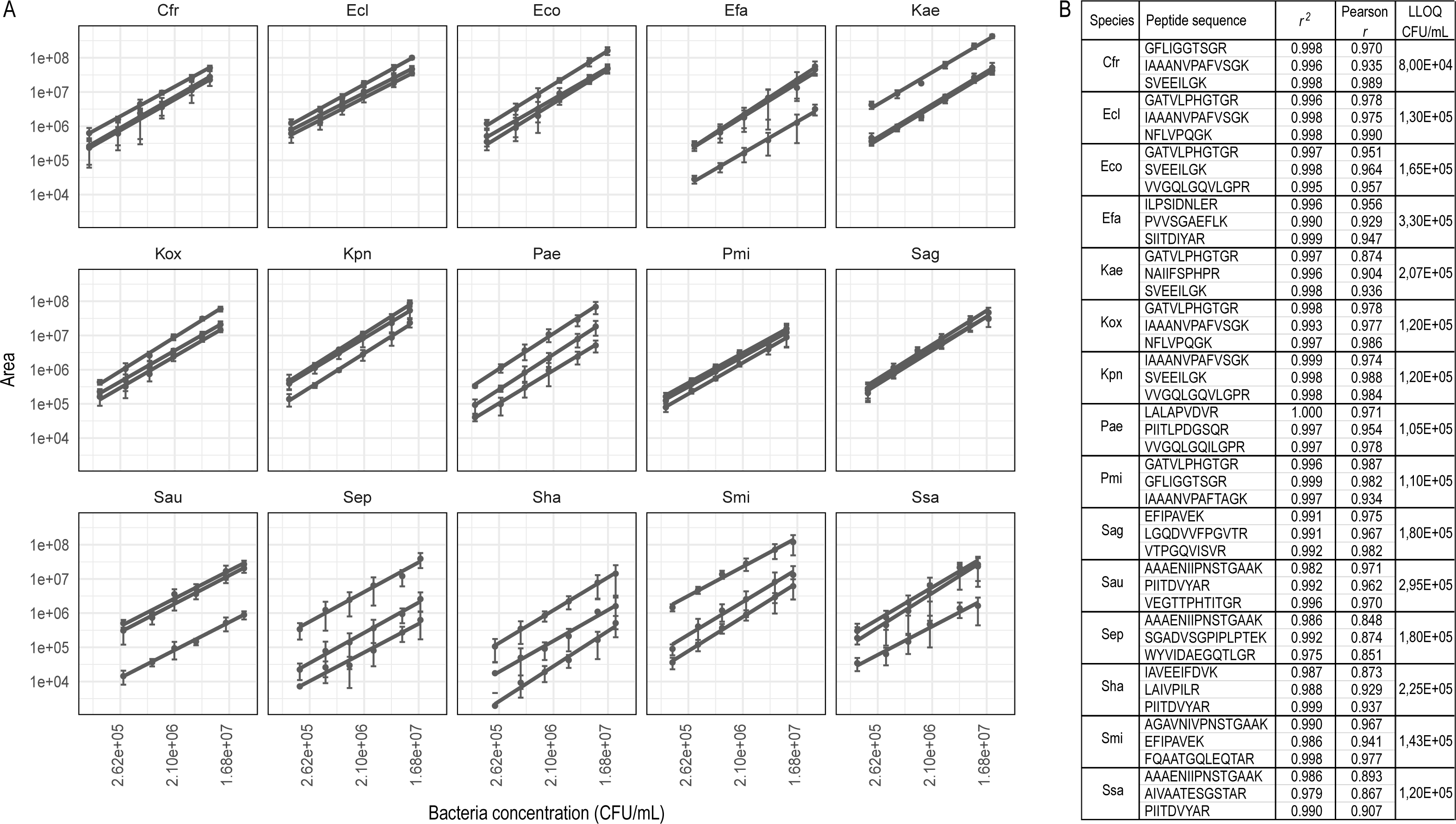
Linear regressions of peptides used for bacterial quantification. A) Linearity curves obtained for the 3 peptides used to quantify each of the 15 species (quantifier peptides) and calculated using the log_10_ of bacterial concentration (CFU/mL) across 6 different points and the log_2_ of the area of the quantifier peptides. B) Corresponding coefficients of determination (r^2^), Pearson correlation coefficient (r) and LLOQ for the 3 quantifier peptides.

#### Reproducibility of quantification

The reproducibility of quantification of the LC-SRM-ML workflow was then evaluated at 3 levels: analytical (LC-SRM injections), technical (whole workflow), and biological (whole workflow with different urines). To do so, the four more prevalent species causing UTI (*E.coli* and *K. pneumoniae*, Gram-negative species and *E. faecalis* and *S. agalactiae*, Gram-positive species) were spiked at a concentration > 1×10^6^ CFU/mL (Supplemental Table S1) in 2 different backgrounds (50 mM Tris or urine), in 6 replicates. The Tris buffer background was used to evaluate the technical reproducibility, from the sample preparation to the injection in LC-SRM, independently of the effect of the biological matrix. As only a half of the sample was injected, the other half of the samples prepared in the Tris buffer background were pooled together and injected 6 times to measure the reproducibility of the LC-SRM system. The effect of the urine matrix on the analysis (biological reproducibility) was evaluated with bacterial inoculations in urine from healthy donors or from UTI-negative (confirmed by bacterial culture) sick patients (specimens collected at the clinical microbiology lab). As urine collected from patients having a bacterial infection and / or having other types of disorders may contains precipitates or hemoglobins and was more turbid than the one from healthy donors, we added a very short centrifuge step at low speed (1 min at 100x*g*) before bacteria isolation, to remove large human cell debris that may lead to technical issue on the LC-MS system.

The signal from the 3 previously defined quantifier peptides was extracted in Skyline for the 4 species and used to calculate the relative standard deviation (RSD (%)) (Supplemental Table S8). As shown in figure 4, an average of 8.6% and 27.1% of variation was measured at the analytical and at technical level respectively. As expected, the quantification reproducibility is reduced for biological samples: 41.9% and 46.4% of RSD were obtained when bacteria were inoculated in urine from healthy volunteers or from patients respectively. We can also notice the reproducibility is slightly improved by adding the rapid centrifugation at the beginning of the process to remove large urine particles, with 42.0% of RSD. More importantly, this step greatly preserves the robustness of the LC-MS system, without compromising the level of bacterial detection as shown by the peptide intensities (Supplemental Figure S4).

**Figure 4:**
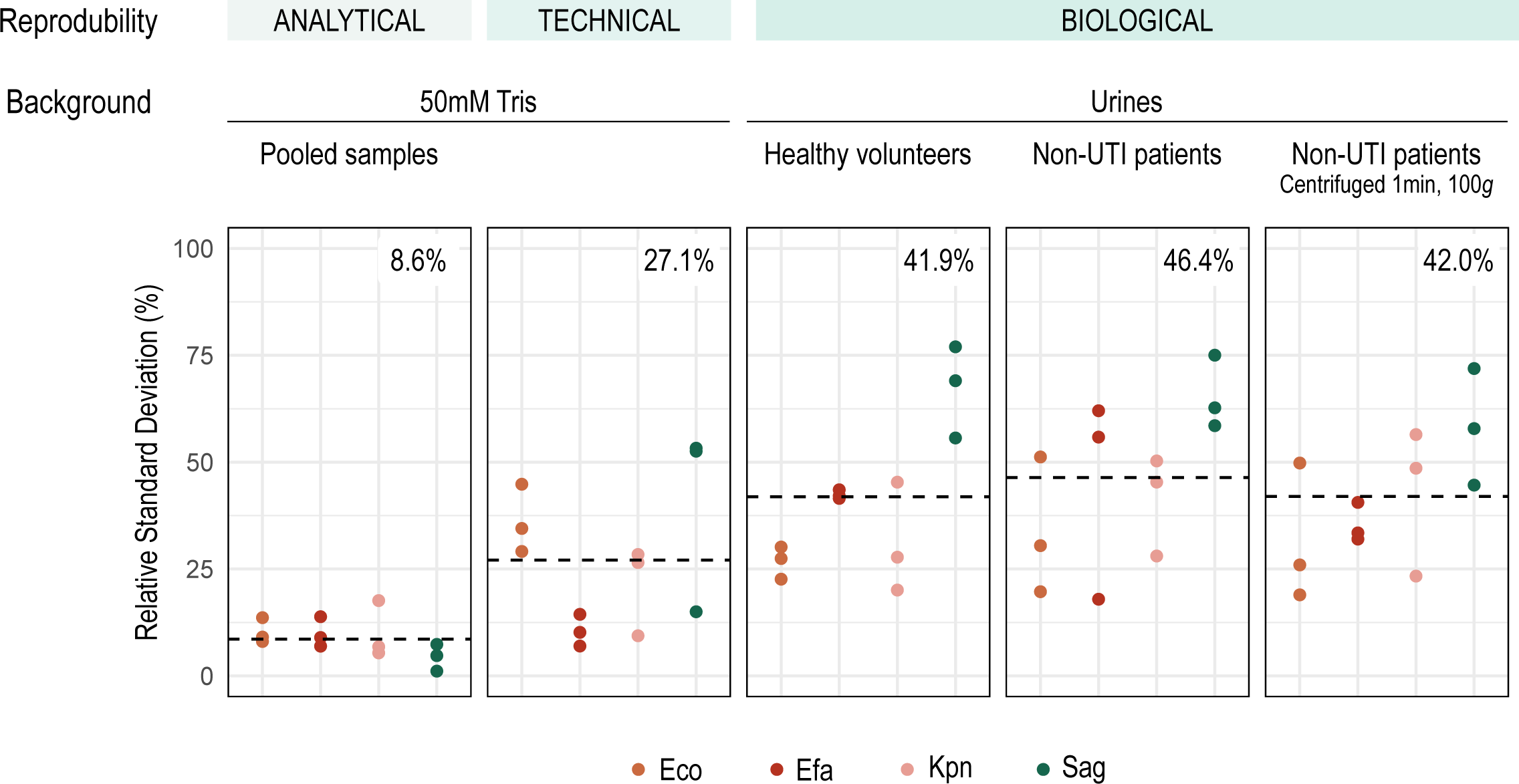
Reproducibility of quantification. Bacterial species Eco and Kpn (Gram+), Efa and Sag (Gram-) were spiked into 2 backgrounds (50 mM Tris or urine), in 6 replicates. Analytical and technical variabilities were evaluated with the tris background while the biological variation was estimated with urine background. The RSD was calculated across the 6 replicates of the area of the 3 quantifier peptides for each bacteria. The dashed line corresponds to the average of RSD for the 4 bacterial species and the corresponding value is indicated at the upper right corner.

### LC-SRM-ML workflow validation

To fully validate our LC-SRM-ML workflow, we collected 84 UTI-positive specimens from patients and compared the results obtained in terms of identification and quantification with the gold standard culture-MALDI. However, due to the unequal prevalence of each bacterial species (43, 44), we obtained samples for only seven species while our method allows the identification of 15 different species. Moreover, the culture-MALDI procedure does not offer a true quantification but only a rough range of infection through a visual estimate of the number of colonies on the culture plate based on the technician’s expertise. In this context, our method cannot be fully validated based on patient specimens. Therefore, we also used a new set of 45 controlled inoculations that were used to compare both methods.

#### Validation on inoculated samples

The 15 bacteria species included in our model were inoculated at 3 different concentrations in healthy urine (Supplemental Table S1) and were analyzed with both the gold standard culture - MALDI procedure (turnaround time 24h) and the LC-SRM-ML (turnaround time 4h) workflow. A plate counting was also performed on the broth culture used for inoculation in order to obtain a quantification value (Figure 5A). As shown on the upper panel of figure 5B, 97.8% and 93.3% of the 45 analyzed specimens are correctly identified by the culture-MALDI and the LC-SRM-ML methods respectively (Supplemental Table S9). While the MALDI confuses two *Citrobacte*r (*freundii* identified as *werkmanii*), a confusion is made on two *Klebsiella* inoculations *(pneumoniae* predicted as *aerogenes*) with the LC-SRM-ML method. Also, one low concentration (2.8×10^5^ CFU/mL) *S. aureus* inoculation was not classified (*i.e* predicted as “Unknown”) probably due to the low number (4) of detected peptides. Nevertheless, in terms of quantification, the LC-SRM-ML workflow outperforms the rough estimation of the bacterial concentration given by the culture-MALDI method (Figure 5B, bottom panel). In 15.5% of the cases, the range (1×10^5^ - 1×10^5^ CFU/mL) given is out of the expected concentrations, resulting in an underestimation of the bacterial infection. An accurate quantification could be achieved with our LC-SRM measurement as demonstrated by the Pearson correlation factor of 0.757 obtained between the plate counting performed on the broth culture and the bacterial concentration calculated based on the LC-SRM signal across the 45 inoculated urines (Figure 5C).

**Figure 5:**
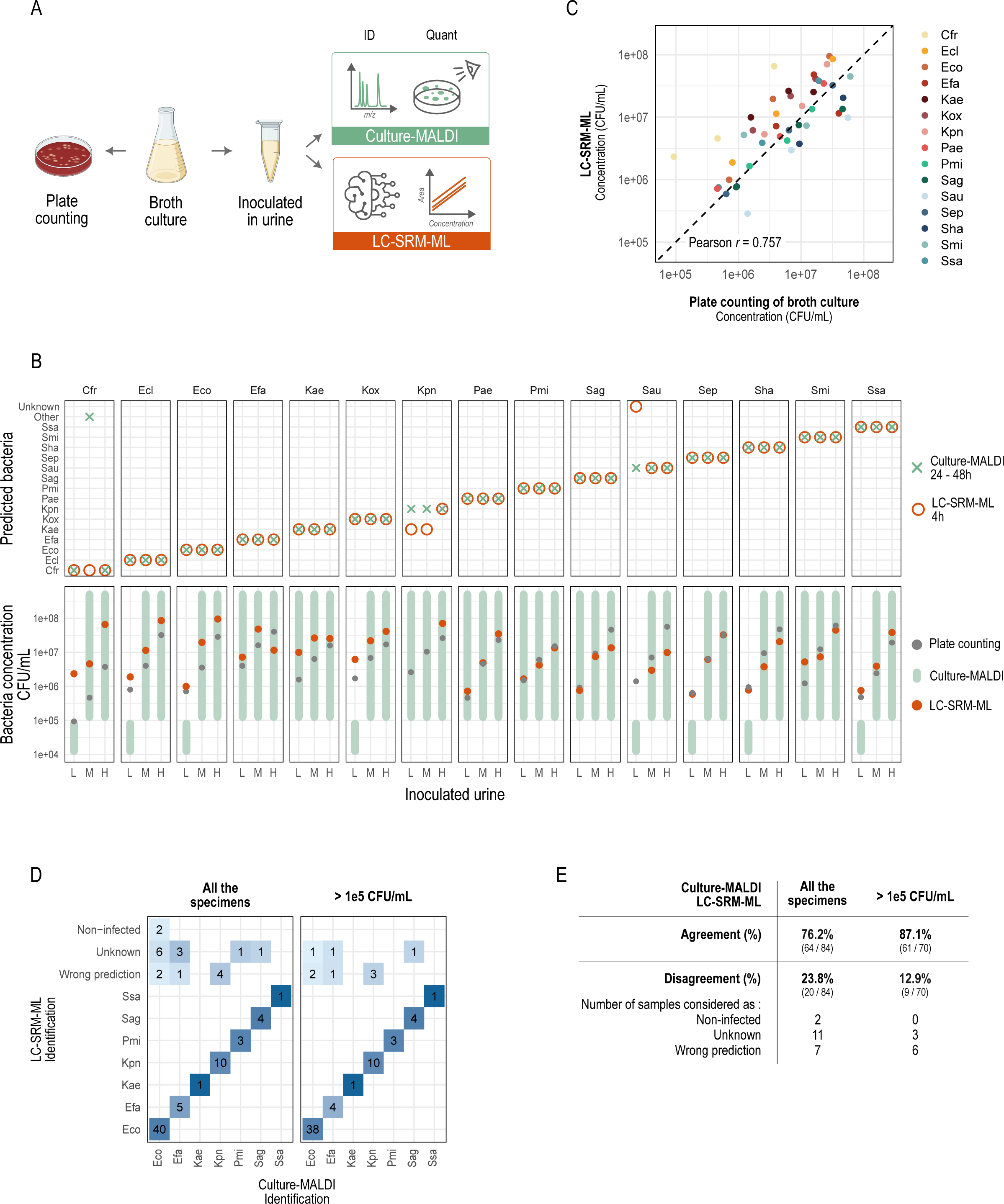
Validation of the LC-SRM-ML workflow. A) Method used for the validation on inoculated urines. B) The 15 bacterial species have been inoculated in urine from healthy volunteers, each at 3 concentrations (L: low; M: medium; H: high) and submitted to the culture-MALDI (green) or the LC-SRM-ML (orange) workflows. Upper and lower panels respectively correspond to the identification and the quantification of the bacterial species in each sample with both methods. C) Pearson correlation between the inoculated bacterial concentration in CFU/mL (calculated from culture plate counting) and the measured bacterial concentration with the LC-SRM-ML method in CFU/mL, for the 45 inoculated urines. D) Identifications of 84 UTI-positive specimens collected at the CHU de Québec, for all the samples (left panel) or for high bacterial level (> 1×10^5^ CFU/mL) (right panel). E) Numbers of urines and corresponding percentage (indicated in bold) of identification agreement or disagreement between the LC-SRM-ML method and the gold standard culture-MALDI, over 84 UTI-positive specimens.

#### Validation on urines from UTI-positive patients

Among the 84 patients specimens tested, 76.2% of the predictions given by the LC-SRM-ML method were in agreement with the culture-MALDI gold standard procedure (Figure 5D and 5E, Supplemental Table S10). The concordance reached 87.1% (61 samples over 70) when looking at high bacteria content level (> 1×10^5^ CFU/mL, determined with the culture-MALDI) with a hesitation of identification between two species (same probabilities) for 5 specimens. At this level, the machine learning prediction model was not able to assign a bacteria identification for 3 specimens (only 4 to 7 peptides of the UTI signature detected), and an error of prediction was performed for 6 specimens. However, even though the culture-MALDI is the gold standard in the clinical laboratories to analyze specimens, we and others already have reported errors of uropathogens identification on inoculated samples (29, 45) with this method. We can also notice that the quantification calculated with the LC-SRM signal falls out of the range estimated with the culture-MALDI for 3 specimens (Supplemental Figure S5), while for the 61 others, we obtained a precise measurement rather than a range.

## DISCUSSION

In the context of the emergence of multi drug resistant pathogens (46–48), the rapid and reliable identification of microbial species is becoming essential, in order to provide appropriate antibiotherapy to patients and to reduce the occurrence of new antimicrobial resistances (49). To date, MALDI-TOF analysis is the gold standard in clinical laboratories for urine analysis but the long bacterial culture step (24h-48h) required to obtain pure colonies prior to the analysis can delay appropriate treatments.

Here, we have optimized a fast and robust workflow, called LC-SRM-ML, combining targeted mass spectrometry and machine learning-defined peptide signature, to accurately identify and quantify uropathogens, without the need for bacterial culture. With this method, the 15 most prevalent bacterial species, representing 84% of all UTIs, could be distinguished with a signature of 82 peptides. Compared with our first version of the method on orbitrap instruments, this workflow can be operated from just 1mL of urine (previously 10 mL) and on triple quadrupole mass spectrometers, whose cost and ease of use are well suited to clinical microbiology laboratories. The workflow relies on a short and automatable sample preparation and a fast LC gradient (10 min), enabling identification and quantification of the species responsible for the infection in less than 4 hours and for up to 75 samples analyzed per day.

Our results demonstrated that the LC-SRM-ML workflow can correctly identify over 87% of the samples. However, 3 *Klebsiella* species included in the model share 21 peptides of the UTI signature, making them difficult to distinguish. To obtain a better classification, further experiments could be done to enrich the UTI signature with peptides specific to these species. Moreover, we observed that prediction accuracy was lower on clinical specimens, probably due to the wide range of clinical strains that can be found in biological samples (50, 51). It would therefore be useful, as a next step, to include clinical isolates to train a new machine learning model, which would capture the whole biological diversity of the microorganisms. Although these results seem identical to or even slightly below those obtained with MALDI-TOF, it should be remembered that they are obtained with an analysis lasting just 4 hours and without bacterial culture, which is a significant improvement that could eventually enable the physician to prescribe a more targeted treatment, more quickly and only when necessary. In terms of quantification, we showed that our workflow outperformed the estimated range given with the standard method by the visual inspection of the number of colonies on the plate before the MALDI-TOF analysis. In particular, we demonstrated that this method underestimates the level of infection in approximately 15% of cases. This may also be explained by the fact that culture-based methods only count bacterial cells that are still viable, which is not the case with our strategy. However, accurate quantification of bacterial content is important in the case of UTI as it is necessary to distinguish simple contamination of urine from a true infection requiring antibiotherapy (> 1×10^5^ CFU/mL) (18, 52). Moreover, clinical signs do not always correlate with the severity of the infection. Therefore, quantification can provide additional information to the clinician for a better use of antibiotics.

Sample preparation, although already fast, could also be improved in the future on several points. Currently, two hours are dedicated to enzymatic digestion (mutanolysin for protein extraction and trypsin for peptide digestion). Nevertheless, several studies have shown that this time can be reduced to a few minutes without compromising the sensitivity of the analysis (53, 54). Furthermore, as the variability of clinical specimens is relatively high, transferring sample preparation to a liquid handler could be particularly useful to minimize technical variability as well as increasing analytical throughput. This high variability on clinical samples can be explained by the nature of urine. Indeed, we have observed that some of them produce retention time shifts that are also seen on the CytoC internal standard peptides. However, the choice of monitoring each peptide all along the gradient, rather than in a SRM scheduled method, prevents the loss of signal linked to this matrix effect and increases the robustness of our workflow.

While the LC-SRM-ML method still needs further refinement and extensive validation (various strains, double infections, transferability on other triple quadrupole instrument…), before being ready for clinical pre-implementation, the work presented here represents an important proof-of-concept demonstrating that LC-MS in SRM mode combined with artificial intelligence can be a powerful tool for detecting and quantifying bacterial species in biological fluids. The method could indeed be extended to other types of clinical specimens, for example on cerebrospinal fluid in meningitis, from positive blood culture in sepsis or even to detect upper and lower respiratory tract infections (55–57). For UTIs, the LC-SRM-ML workflow could be a first step, used to complement the standard MALDI-TOF procedure as a rapid method for identifying and quantifying the 15 most frequent species only, but it could also be extended to a larger number of clinically relevant species in a later stage, through the development of new machine learning signatures. Moreover, it would be particularly interesting to enrich the peptide signature for the detection of virulence factors (58–61) or antimicrobial susceptibility (62, 63). Whatever the case, it is now clear, confirmed by our study and others (64, 65), that LC-MS and artificial intelligence are methods destined to play a key role in microbial diagnostics in the near future.

## ACKNOWLEDGMENTS

This work was funded by a grant from Génome Québec (GQ130685). The authors acknowledge the instrumental support provided by Thermo Fisher Scientific (San Jose, USA).

## DATA AVAILABILITY

The raw mass spectrometry data are available on ProteomeXchange, on the following identifier: PXD052699. R code is available at https://github.com/clarissegotti/UTI_LC-SRM-ML

## DECLARATION OF INTEREST

Cristina C. Jacob, Claudia Martins and Neloni R. Wijeratne are employees of Thermo Fisher Scientific.

## SUPPLEMENTAL DATA

This article contains supplemental data: supplemental figures S1-S5 and supplemental tables S1-S10.

## CREDIT AUTHORSHIP CONTRIBUTION STATEMENT

**Clarisse Gotti**: Conceptualization, Methodology, Validation, Formal analysis, Investigation, Writing - Original Draft, Visualization. **Florence Roux-Dalvai**: Conceptualization, Methodology, Writing - Original Draft, Supervision, Project administration. **Ève Bérubé**: Investigation, Resources. **Antoine Lacombe-Rastoll**: Investigation. **Mickaël Leclercq**: Software, Writing - Review & Editing. **Cristina Jacob**: Resources. **Maurice Boissinot**: Resources. **Claudia Martins**: Resources, Funding acquisition. **Neloni Wijeratne**: Resources, Funding acquisition. **Michel G. Bergeron**: Resources, Funding acquisition. **Arnaud Droit**: Writing - Review & Editing, Supervision, Project administration, Funding acquisition.

## ABBREVIATIONS

ACN: Acetonitrile
AUC: Area under curve
CE: Collision energy
Cfr: *Citrobacter freundii*
CFU: Colony forming units
DDA: Data-dependent acquisition
DIA: Data-independent acquisition
DTT: Dithiothreitol
Ecl: *Enterobacter cloacae*
Eco: *Escherichia coli*
Efa: *Enterococcus faecalis*
FA: Formic acid
Kae: *Klebsiella aerogenes*
Kox: *Klebsiella oxytoca*
Kpn: *Klebsiella pneumonia*
LC: Liquid chromatography
LLOQ: Lower limit of quantification
MALDI-TOF: Matrix-assisted laser desorption ionization–time of flight
MCC: Matthews Correlation Coefficient
MRM: Multiple reaction monitoring
MS: Mass spectrometry
Pae: *Pseudomonas aeruginosa*
PCR: Polymerase chain reaction
Pmi: *Proteus mirabilis*
RSD: Relative standard deviation
S/N: Signal to noise
Sag: *Streptococcus agalactiae*
Sau: *Staphylococcus aureus*
SDC: Sodium deoxycholate
Sep: *Staphylococcus epidermidis*
Sha: *Staphylococcus haemolyticus*
Smi: *Streptococcus mitis*
SRM: Selected reaction monitoring
Ssa: *Staphylococcus saprophyticus*
UTI: Urinary tract infection
WGS: Whole genome sequencing

