## Supplemental Figure S1 - S5; Supplemental Table S2 for "LC-SRM combined with machine learning enables fast identification and quantification of bacterial pathogens in urinary tract infections"

### Table of contents

#### List of figures


**Supplemental Figure S1**
Quality control — Page S-1

**Supplemental Figure S2**
Bacteria identification by machine learning on inoculated samples — Page S-2

**Supplemental Figure S3**
Linear regressions for all detected peptides of the 15 bacterial species — Page S-3

**Supplemental Figure S4**
Intensities of the 3 quantifier peptides of 4 bacterial species — Page S-4

**Supplemental Figure S5**
Quantification of UTI-positive patient urines — Page S-5


#### List of tables


**Supplemental Table S1**
Dataset and inoculations information — .xlsx file

**Supplemental Table S2**
Bacterial species culture conditions — Page S-6

**Supplemental Table S3**
SRM acquisition parameters and transition list — .xlsx file

**Supplemental Table S4**
Retention times of the 82 UTI signature peptide and 6 CytoC standards — .xlsx file

**Supplemental Table S5**
Quantification data for the *Transfer – Inoculations* dataset — .xlsx file

**Supplemental Table S6**
Linear regressions information for all the detected peptides for each bacterial species for the *Transfer – Inoculations* dataset — .xlsx file

**Supplemental Table S7**
Linear regressions, Pearson correlation coefficient, S/N ratio and LOQ for the 3 quantifier peptides per bacterial species for the *Transfer – Inoculations* dataset — .xlsx file

| | |
|---|---|
| **Supplemental Table S8**<br>Quantification data for the *Transfer* – *Reproducibility* dataset | .xlsx file |
| **Supplemental Table S9**<br>Detailed results for the *Validation* – *Inoculations* dataset | .xlsx file |
| **Supplemental Table S10**<br>Detailed results for the *Validation* – *Patients* dataset | .xlsx file |

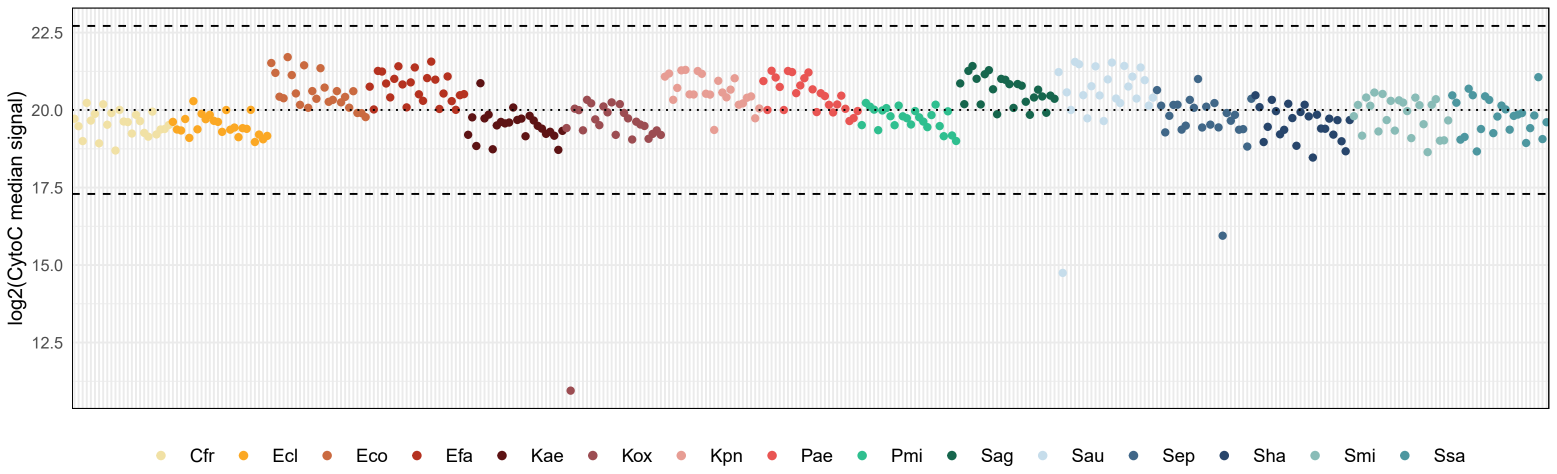


**Supplemental Figure S1 : Quality control**

Injections were controlled with CytoC standard peptides spiked in the sample just before injection. Average log2 area (dotted line) and +/- 3 standard deviation (dashed lines) were calculated over the 360 samples of the Transfer - Inoculations dataset.

A

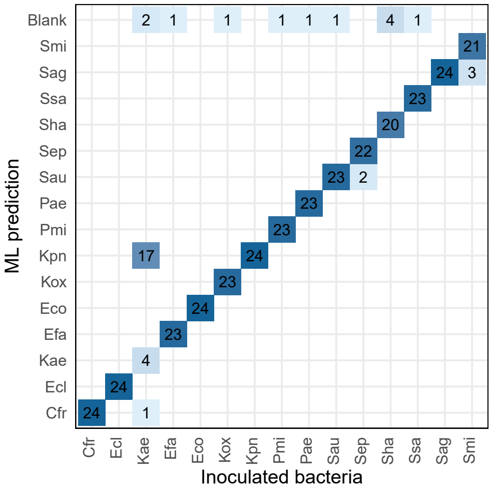


B

| | DIA | | SRM | |
|---|---|---|---|---|
| | Kae | Kpn | Kae | Kpn |
| SVEEILGK | 0 | 8 | 24 | 24 |
| STAETIVYSALETLAQR | 0 | 8 | 18 | 24 |
| AQYVLAEQVTR | 0 | 7 | 13 | 23 |
| FGGESVLAGSIIVR | 10 | 0 | 0 | 0 |
| VASLEGDVLGSYQHGAR | 12 | 0 | 0 | 0 |

**Supplemental Figure S2 : Bacteria identification by machine learning on inoculated samples**

A) Confusion matrix of bacteria inoculated in healthy urines. Predictions by machine learning were done with the first generation model obtained from DIA data. B) Comparison of specific peptide signature to distinguish K. aerogenes (Kae) and K. pneumoniae (Kpn), in DIA and SRM data.

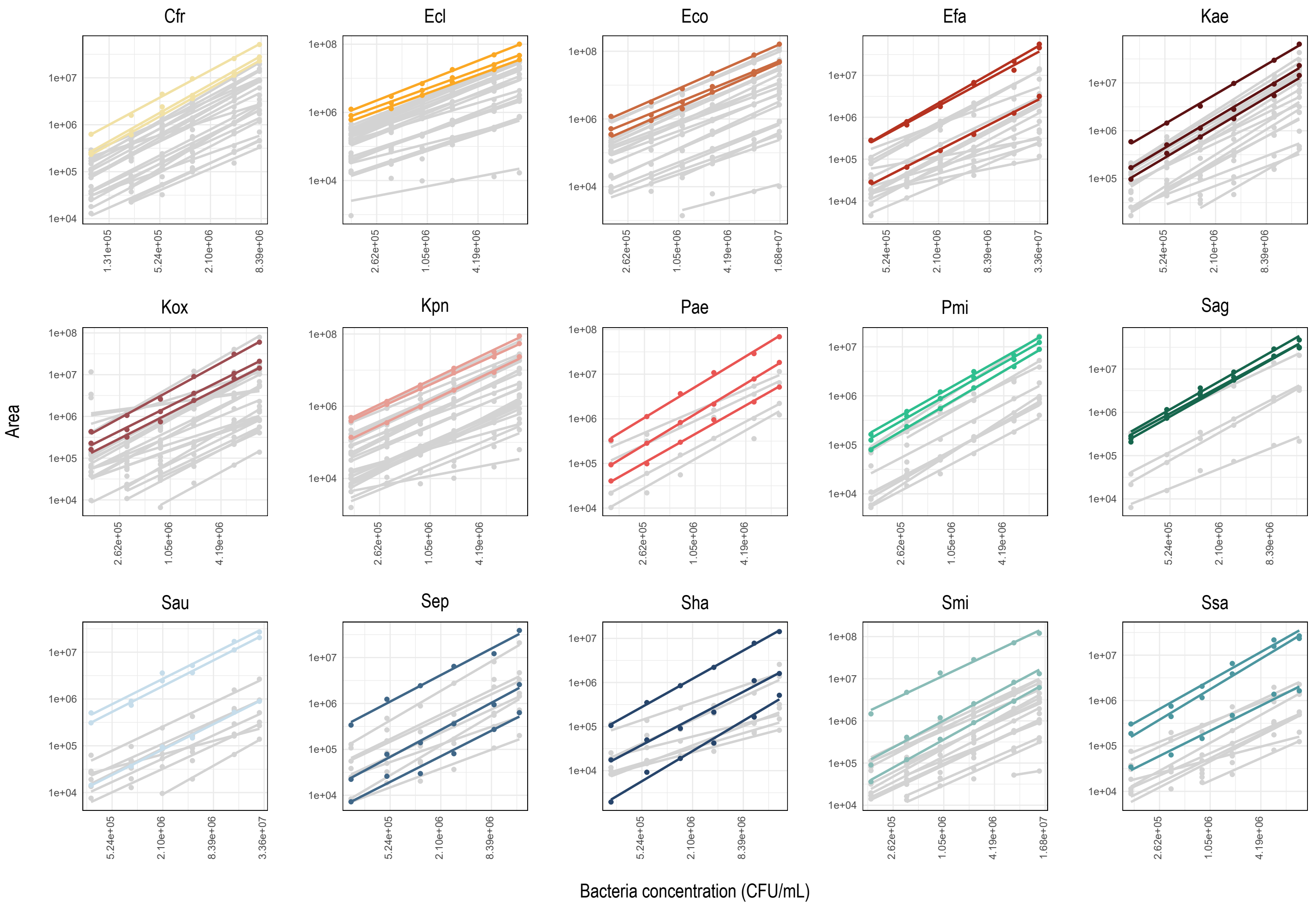


**Supplemental Figure S3 : Linear regressions for all detected peptides of the 15 bacterial species**
Linear regressions were calculated for all the detected peptides for each bacterial species, between the $log_2$ of the peptide peak areas and the $log_{10}$ of the bacterial concentration inoculated in urine (CFU/mL). Peptides in colors are those selected for the quantification of unknown samples while those in grey (+ colored) will be only used for bacterial prediction by machine learning.

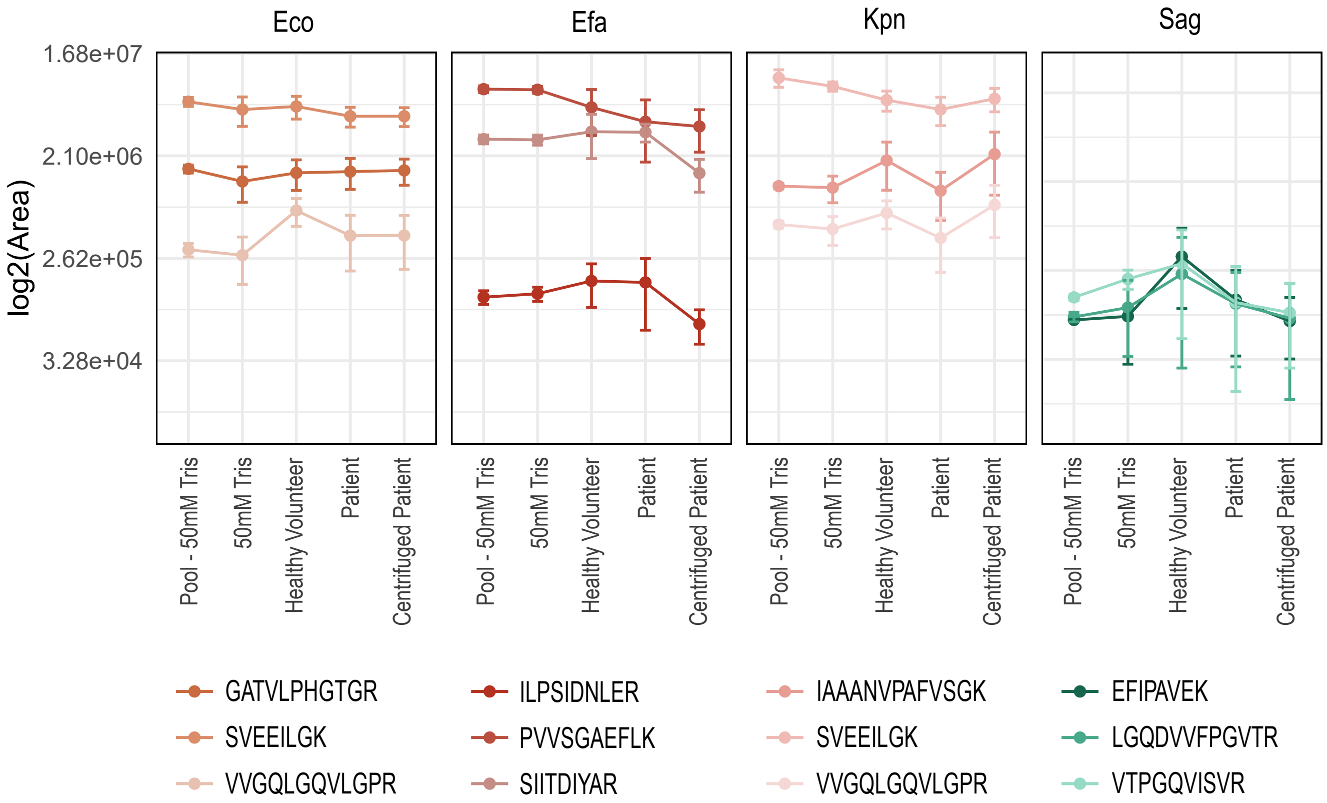


**Supplemental Figure S4 : Intensities of the 3 quantifier peptides of 4 bacterial species**

The $log_2$(Area) of each quantifier peptide was plotted for *E. coli* (Eco), *E. faecalis* (Efa), *K. pneumoniae* (Kpn) and *S. agalactiae* (Sag) for the 5 reproducibility testing methods : Pool - 50 mM Tris (injection); 50 mM Tris (technical); urines from healthy volunteers or from patients with or without a small centrifugation before bacterial isolation (biological).

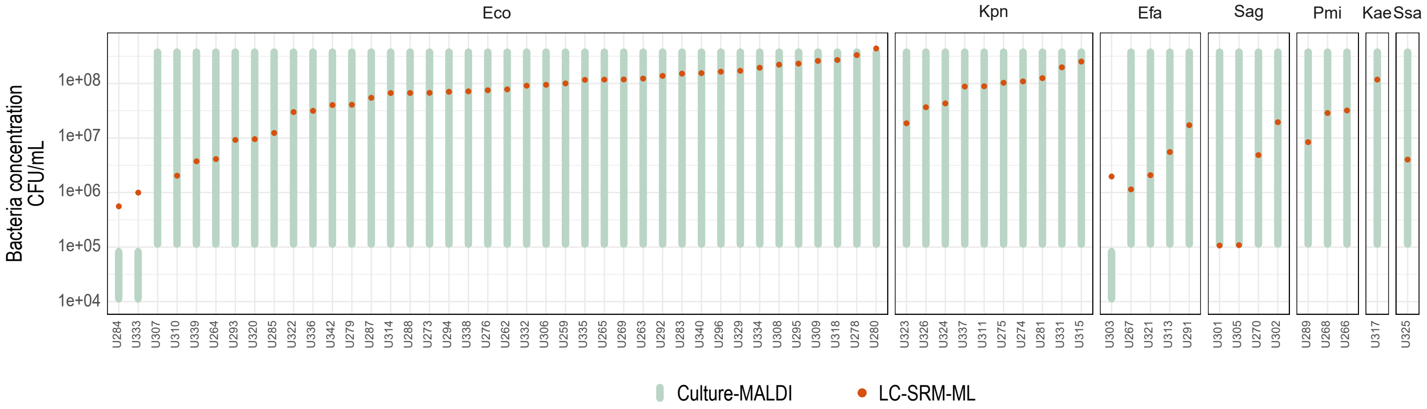


**Supplemental Figure S5 : Quantification of UTI-positive patient urines**

The 64 UTI-positive urines from patients whose machine learning prediction was in agreement with MALDI-TOF analysis were quantified by plate counting performed before MALDI-TOF analysis (green) or by LC-SRM (orange).

| Bacterial species | Abbreviation | Strain | Culture condition | Gram-stain | Incubation time | Semi-log culture concentration (CFU/mL) |
|---|---|---|---|---|---|---|
| *Citrobacter freundii* | Cfr | CCRI-429 | aerobe | negative | 2h10 | 2,20E+08 |
| *Enterobacter cloacae* | Ecl | CCRI-860 | aerobe | negative | 1h30 | 2,00E+08 |
| *Escherichia coli* | Eco | CCRI-12923 | aerobe | negative | 1h50 | 4,70E+08 |
| *Enterococcus faecalis* | Efa | CCRI-95 | aerobe | positive | 3h00 | 1,10E+09 |
| *Klebsiella aerogenes* | Kae | CCRI-446 | aerobe | negative | 1h40 | 5,40E+08 |
| *Klebsiella oxytoca* | Kox | CCRI-1109 | aerobe | negative | 2h00 | 5,20E+08 |
| *Klebsiella pneumoniae* | Kpn | CCRI-563 | aerobe | negative | 1h40 | 3,90E+08 |
| *Pseudomonas aeruginosa* | Pae | CCRI-691 | aerobe | negative | 3h00 | 9,30E+07 |
| *Proteus mirabilis* | Pmi | CCRI-675 | aerobe | negative | 1h40 | 4,40E+08 |
| *Streptococcus agalactiae* | Sag | CCRI-235 | air + 5% $CO_2$ | positive | 2h30 | 2,30E+08 |
| *Staphylococcus aureus* | Sau | CCRI-9443 | aerobe | positive | 1h50 | 3,10E+08 |
| *Staphylococcus epidermidis* | Sep | CCRI-203 | aerobe | positive | 2h40 | 9,60E+07 |
| *Staphylococcus haemolyticus* | Sha | CCRI-210 | aerobe | positive | 3h10 | 1,40E+08 |
| *Streptococcus mitis* | Smi | CCRI-259 | air + 5% $CO_2$ | positive | 4h45 | 1,30E+08 |
| *Staphylococcus saprophyticus* | Ssa | CCRI-223 | aerobe | positive | 2h50 | 2,10E+08 |

**Supplemental Table S2 : Bacterial species culture conditions**
